# Shift augmentation improves DNA convolutional neural network indel effect predictions

**DOI:** 10.1101/2025.04.07.647656

**Authors:** Anya Korsakova, Divyanshi Srivastava, David R Kelley

**Affiliations:** Calico Life Sciences, South San Francisco, CA 94080, USA

## Abstract

Determining genetic variant effects on molecular phenotypes like gene expression is a task of paramount importance to medical genetics. DNA convolutional neural networks (CNNs) attain state-of-the-art performance at predicting variant effects on gene regulation. However, most applications of such models focus on single nucleotide polymorphisms (SNPs), as technical challenges limit their application to insertions and deletions (indels). Sequence shifts from indels introduce technical variance in deep CNNs through misalignment of pooling blocks and output boundaries, creating artificially inflated variant effect scores compared to SNPs and confounding their interpretation. In this work, we demonstrate this technical variance in model predictions and present two strategies based on data augmentation with sequence shifts that reduce it. Applied to the state-of-the-art Borzoi model, our stitching approach improves indel eQTL classification accuracy across GTEx tissues. Furthermore, we demonstrate these techniques and observe compelling eQTL concordance for larger structural variants and tandem repeats. We additionally introduce *in silico* deletion (ISD) as an interpretation technique and validate it using MPRA data, demonstrating concordance between predicted and experimental measurements for deletion effects. Our strategies expand the utility of regulatory sequence machine learning for studying the full spectrum of noncoding genetic variation in human development and disease.

## 1 Introduction

Dissecting and precisely understanding the contribution of DNA sequence to cell type-specific regulatory activity is a key ongoing challenge of genome research. Deep learning methods have been successfully applied across diverse regulatory stages, achieving state-of-the-art accuracy at predicting various measurements on unseen DNA sequences, including gene expression [1, 2], RNA splicing [3, 4], alternative polyadenylation [5, 2], transcription initiation [6, 7, 8], RNA degradation rate [9], transcription factor (TF) binding [10, 11], and 3D genome contacts [12, 13]. Deciphering the consequences of inherited and acquired mutations is a particularly important application of such methods. Neural networks are capable of rapidly predicting the regulatory effects of individual variants via comparing reference versus alternative sequence predictions. These variant effect predictions may be used to study rare and de novo variants in disease [14] and prioritize putative causal variants from GWAS [15].

Most genetic variant analyses thus far have focused on substitution single nucleotide polymorphisms (SNPs), while insertions and deletions (indels) have received less attention. However, small indels make up about 24% of common variants in the gnomAD v4 database [16] and are equally enriched for heritability compared to SNPs [17]. Thus, ignoring indels in genetic analyses risks missing valuable insights and incorrectly prioritizing SNPs.

In our experience developing regulatory sequence deep learning models, we find that estimating indel effects has technical challenges. These models take a fixed length one-hot-encoded nucleotide sequence as input. When an indel is introduced, the sequence shifts on at least one side of the alteration. As we will show below, these shifts produce undesirable technical variance (i.e. nonbiological) due to changing boundaries of the predicted output and convolution blocks. In this work, we explain this problem and measure its influence on estimated indel effect scores. Although technical variance in indel effect estimation cannot be completely absolved in our current framework, we propose and evaluate techniques to alleviate it. We further demonstrate how these strategies expand the applicability of a large multi-task regulatory sequence model Borzoi, an extension of the Enformer model [1] to RNA-seq, to a broader class of tasks involving comparisons between inserted or deleted alternative alleles.

## 2 Results

### 2.1 Boundary shifts introduce technical variance in DNA convolutional networks

Deep neural networks for DNA sequences usually consist of repeated blocks that include convolution, normalization, activation, and pooling operations. The pooling operation typically computes an unparameterized function of a local window, e.g. taking the channel-wise maximum. Input sequence shifts (that are not multiples of the window size) change pooling window boundaries, producing different values for the downstream computations (Fig. 1A). The model outputs, whether they represent a single prediction or a sequence of predictions, correspond to specific boundaries, which also shift.

**Figure 1:**
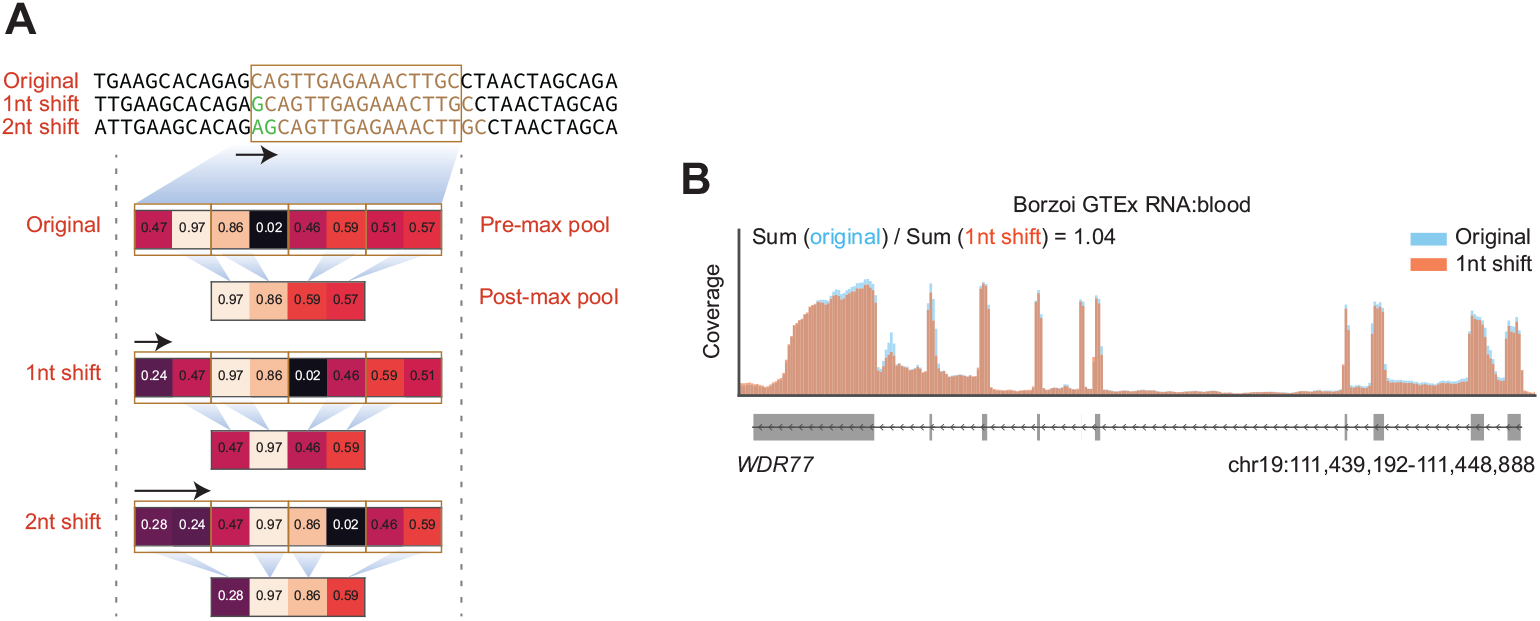
Sequence shift effect on pooling boundaries and predictions. A) Shifting an example sequence before a max pooling block affects the boundaries of the operation. For a width 2 max pool, the 2 nt shift output is similar, but with all values shifted by one position. However, the 1 nt shift changes some output values because the max operation is computed between different adjacent pairs. B) Shifted sequences may produce considerably different predictions. For the *WDR77* gene sequence, Borzoi predicts blood RNA-seq coverage with fold change of 1.04 upon shifting the sequence 1 nt to the left. Coverage from three GTEx:blood RNA-seq tracks were averaged for the calculation and plot.

We analyzed such shifts using Borzoi, a recent model trained to predict RNA-seq, CAGE, DNase/ATAC-seq, and ChIP-seq coverage in 32 bp bins across a 524 kb sequence. A 1 nt shift of the sequence containing the *WDR77* gene exemplifies the problem, producing a 1.04x fold change in predicted blood RNA-seq coverage, despite representing the same DNA sequence (Fig. 1B). Many expression QTLs (eQTLs) have effect sizes within this range, highlighting that distinguishing a true indel eQTL from this background shift variance would be difficult. Larger indel shifts would be expected to produce even more substantial changes to the model predictions, interfering with the downstream interpretation of model scores.

### 2.2 Indel effect predictions skew towards greater magnitudes

To predict genetic variant effects using deep learning, typically we center the input sequence on the variant. Insertions and deletions inevitably introduce input sequence shifts like the one above. A baseline insertion procedure might insert the additional nucleotides in place, shift the adjacent rightside nucleotides rightward, and trim the far right nucleotides to maintain a fixed length (Fig. 2A). A baseline deletion procedure might delete the nucleotide, shift the adjacent right-side nucleotides leftward, and pad the far right of the sequence to maintain a fixed length. Both scenarios produce misaligned nucleotides (and thus pooling and output boundaries) for half the sequence. In addition to the shifts, insertions lose some sequence outside of the input region, and deletions add padded null values (or new reference sequence) at the edge. These factors lead to non-biological technical variance in indel effect scores.

**Figure 2:**
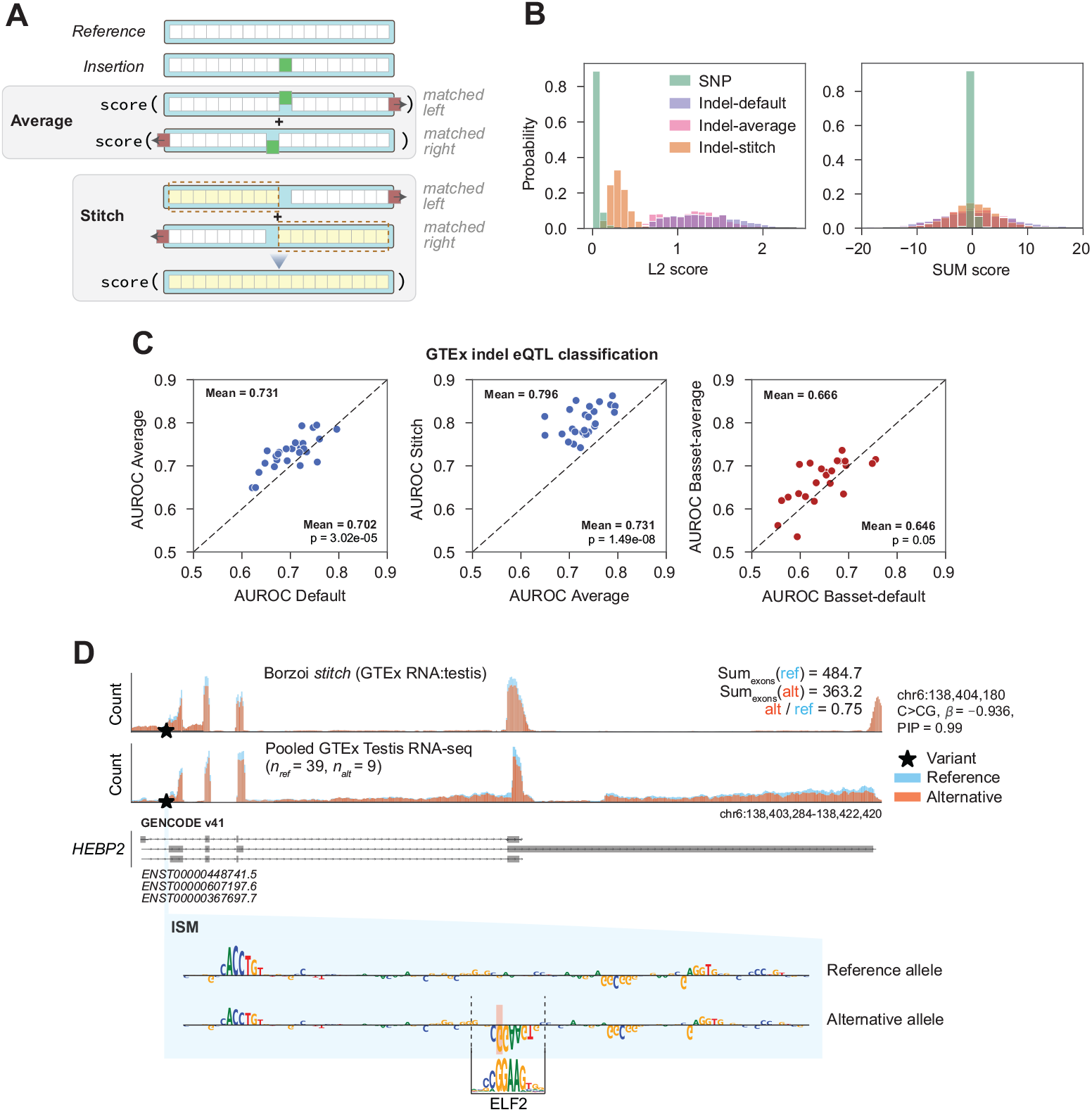
Indel shift augmentation improves effect predictions. A) Insertion strategy depiction. Baseline procedures might match the left side, misalign the right side, and push out some edge sequence on the right, or vice versa. Our “average” strategy averages the scores for the “leftmatched” and “right-matched” procedures. Our stitch strategy concatenates predictions from output bins that remain matched in the left and the right shift procedures, then computes the score for the resulting vector. B) L2 and SUM score distributions for N=3,000 common SNPs and N=3,000 common 1 bp indels with the default (left-matched), average, and stitch strategies. We created a single variant effect score by averaging across all Borzoi output tracks. C) Shift augmentation strategies (average and stitch) improve fine-mapped indel classification performance for random forest classifiers trained on both Borzoi and Basset outputs on 1,168 unique fine-mapped GTEx eQTLs. Each point in the scatterplot represents a GTEx tissue. D) Predicted testis RNA-seq coverage for chr6:138,404,180 C*>*CG eQTL variant in the promoter of the *HEBP2* gene, and measured RNA-seq counts from testis samples pooled over n=39 reference and n=9 heterozygous variant individuals. The eQTL has a PIP=0.99 and *β*_*posterior*_ = −0.936. *In silico* saturation mutagenesis (ISM) maps for the reference and alternative alleles reveal the introduction of a new ETS domain motif with repressive influence.

This technical variance depends on the exact scoring metric used to compare the reference and alternative allele predictions. Regulatory sequence models can be broadly classified into two categories. One, models that predict a single value (such as the presence or absence of a peak) typically score variants by computing the difference or log fold change between reference and alternative allele predictions. Two, models that predict a sequence of values (such as aligned read coverage) across the input sequence compute one of several statistics to compare the pair of vectors and collapse the spatial length axis. These include the sum of the difference or log fold change at each position in the predicted vectors (SUM), L2 norm, or Kullback-Leibler divergence (after normalization) [11]. Here, we focus on SUM scores, which maintain the prediction sign, and L2 norm, which is unsigned, but demonstrates greater sensitivity to complex and subtle effects [2].

To estimate the technical variance in indel scoring, we scored a set of 3,000 SNPs and 3,000 common indels (minor allele frequency ≥ 0.05) in the gnomAD v4 database [16] with Borzoi (see Methods). We focused on 1 bp indels, for which we expect the true effect size distributions to be similar, as one nucleotide is affected in both the indels and SNPs. Instead, the predicted variant effect distributions drastically differed between SNPs and indels for both the SUM and L2 scores (Fig. 2B). While both the SNP and indel SUM scores are centered at zero, the indel distribution has significantly greater variance (“indel-default” in Fig. 2B, *σ*^2^ = 77.07, vs. SNP *σ*^2^ = 0.31). The SNP L2 distribution has a roughly exponential shape, peaked near zero, indicating that most variants alter the coverage track predictions negligibly, but some can have large effects. In contrast, the indel L2 distribution shifts to greater magnitudes (*µ* = 1.34 for “indel-default” in Fig. 2B vs. *µ* = 0.05 for SNP), reflecting the technical variation from the factors described above. This makes joint analysis in which SNP and indel effect predictions must be compared challenging because even nonfunctional indels have greater magnitude scores than most SNPs.

### 2.3 Shift augmentation strategies alleviate variance

Practitioners have at least two equally valid options when introducing indel alleles into the reference sequence. Considering an insertion, one can exactly match nucleotides to the left of the variant, insert the alternative allele, shift the nucleotides to the right of the variant, and truncate at the input sequence length, removing nucleotides from the right edge. Alternatively, one can flip this recipe. In other words, one can introduce the insertion to the right or left of the sequence center. Importantly, the first option has exactly matched pooling block and output boundaries for all nucleotides to the left of the variant (referred to as “left-matched”), and the second option has exactly matched pooling block and output boundaries for all nucleotides to the right of the variant (“right-matched”, Fig. 2A). We will exploit this property to derive variance-minimization strategies. For insertions larger than 1 bp, additional options emerge where one inserts some nucleotides on the left and some on the right, but these options have shifted boundaries on both sides of the variant, making them less interesting for our strategies. Deletions can be handled analogously.

As demonstrated above, shifted boundaries create technical variance. We therefore hypothesized that regions with matched boundaries could be exploited to alleviate variance. Thus, we compute predictions for the reference sequence and then both the left-matched and right-matched alternative sequences. Our first strategy acknowledges the inclusion of misaligned boundary regions, but takes the average of both the left and right shifts to mitigate the effect of newly inserted or deleted nucleotides (“average”). Our second strategy ignores predictions in shifted regions by stitching together left side predictions from the left-matched prediction and right side predictions from the right-matched prediction (“stitch”) (Fig. 2A). The “stitch” recipe is only applicable for models with sequential outputs; models with single value outputs (e.g. peak probabilities) can only use the “average” recipe. For insertions, stitching of the alternative sequences with left- and right-matched shifts should be performed. For deletions, stitching the alternative sequence double counts nucleotides adjacent to the reference, and one should instead introduce shifts and stitch the reference sequence, as if the reference were an insertion relative to the alternative sequence. See the Methods for a more verbose explanation and Supplem. Fig. S1 for examples.

We assessed these strategies on the common SNPs and indels studied above. For tests yielding extremely small P-values (*p* ≤ 1 × 10^−9^), we reported a lower bound of 1 × 10^−9^ to avoid numerical precision issues while still indicating the high level of statistical significance. Indels scored with the average strategy have a slightly lower SUM score variance compared to the default strategy (trimmed Levene’s test *p* ≤ 1 × 10^−9^). Their L2 score distribution is similar to the default strategy, still varying to larger values relative to the SNP L2 score distribution. The stitch strategy mitigated some of the SUM score variance (default vs. stitch variance trimmed Levene’s test *p* ≤ 1 × 10^−9^) and significantly shifted the L2 score distribution towards smaller values compared to the default strategy (two-sided Mann-Whitney U test *p* ≤ 1 × 10^−9^). However, both still differ visibly and significantly from the SNP score distributions (SUM: trimmed Levene’s test *p* ≤ 1 × 10^−9^; L2: two-sided Mann-Whitney U test *p* ≤ 1 × 10^−9^). These observations highlight the importance of the misaligned sequence for indel scoring and point to some improvement alleviating technical variance.

### 2.4 Shift augmentation strategies improves indel eQTL benchmarking

To evaluate the indel scoring strategies, we measured their ability to discriminate fine-mapped indel eQTLs from a negative set of variants. We constructed a benchmark task using 1,168 small indels (≤ 8 nt) fine-mapped to have *>* 0.9 posterior probability of being causal in some tissue in the GTEx project and selected negative indels that match the inserted/deleted nucleotides but lack evidence of being a causal eQTL (Methods). We aimed to predict whether a given indel is a causal eQTL based on Borzoi variant effect predictions using the different scoring strategies provided to a random forest classifier that considers all output tracks. For SNPs, L2 scores outperform SUM scores for this task [2], so we focused on indel L2 scores.

First, we trained tissue-wise classifiers on the left-matched baseline, achieving a mean AUROC of 0.702 across tissues (Fig. 2C). Averaging the left- and right-matched scores, despite still allowing for mis-alignment, increases causal eQTL classification accuracy to AUROC 0.731. By avoiding mis-alignment, stitching the left and right shift predictions further improved the mean AUROC to 0.796.

Fig 2D displays an example fine-mapped insertion eQTL. chr6:138,404,180 C*>*CG attenuates expression of the *HEBP2* gene in testis (*β*_*posterior*_ = −0.936, PIP= 0.99), which is correctly captured by the Borzoi prediction. *In silico* mutagenesis (ISM) revealed the insertion creates a new TF motif that best matches ELF2 in the CISBP2 database [18], but may be bound by similar ETS-domain TFs such as ETV6 or ELF4, for which repression has been reported [19, 20].

While models like Borzoi that predict sequential coverage vectors are especially sensitive to indel boundary shifts, models that predict single values are also affected by indel shifts via their pooling blocks. We additionally benchmarked the DNase peak prediction model Basset against the same eQTL classification task (Methods) [21]. For Basset, the baseline left-matched classifiers achieved AUROC 0.646, and the average shift strategy improved Basset classifiers to mean AUROC 0.666 (Fig. 2C).

We evaluated how well we can predict the effect of indels on the target gene expression, using effect size estimates from the same fine-mapped GTEx eQTL variants. Default SUM score predictions for the tissue-matched tracks achieved an unimpressive Spearman *ρ* = 0.052. The average strategy increases performance to *ρ* = 0.078, and the stitch strategy further increases it to *ρ* = 0.162. Altogether, these results highlight that alleviating the technical noise of indel scoring improves concordance with eQTL statistics.

### 2.5 Shift augmentation stitching extends to larger structural variation

Structural variation (SV) refers to larger insertions, deletions, inversions, and other more complex differences that require more careful genotyping strategies. We hypothesized that our shift mitigation strategies would extend to large indel SVs, too, and enable meaningful Borzoi predictions for these variants. We constructed a benchmark from 9,832 unique marginally associated SV eQTLs from GTEx [22] and a matched set of negative variants. For each positive SV, we sampled a negative non-eQTL SV with closest matched size and at least one expressed gene exon within the model input window (Methods). Despite the noisy positive set due to the absence of fine mapping, the model is able to distinguish between the positive set of eQTL SVs and the negatives with an AUROC 0.588 with the default strategy. Again, we observe that minimizing misaligned bins with the stitch strategy improves AUROC to 0.603 (Fig. 3A). Fig. 3B shows an example of two variants (one SNP, one SV deletion) in perfect LD (*r*^2^ = 1.0) [23] discovered as an eQTL for *CCND3* in GTEX whole blood and associated with blood traits (mean corpuscular hemoglobin and mean corpuscular volume). Only the SV, a 537 nt deletion downstream of the gene, located 44 kb upstream from its TSS, is predicted by Borzoi to alter *CCND3* RNA-seq. ISM attribution scores of the reference allele show multiple regulatory motifs with established relevance in blood cells: *ZNF560, POU3F2/POU3F3/POU3F4*, and *SNAI2* (Fig. 3C).

**Figure 3:**
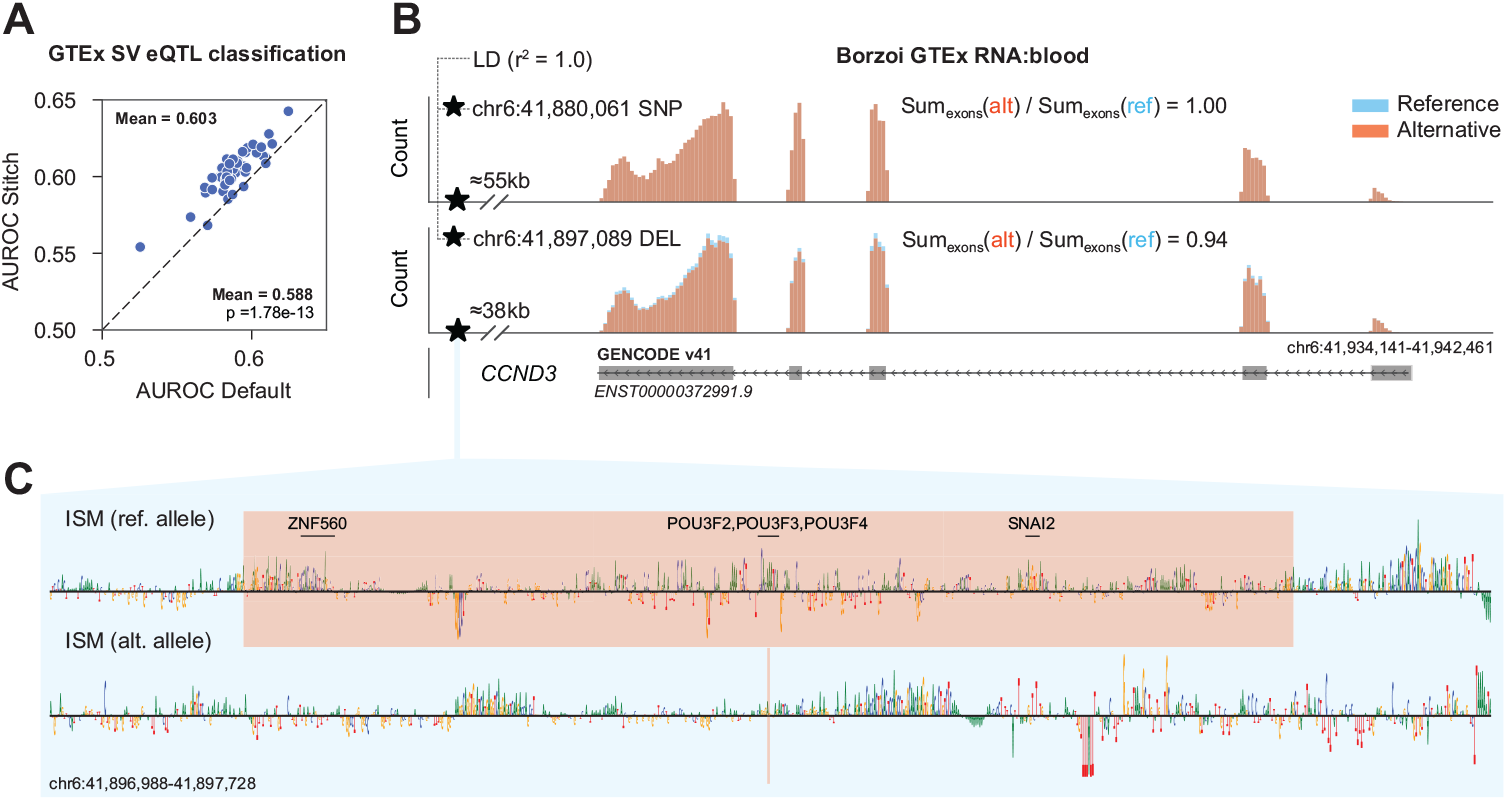
Shift augmentation stitching improves structural variant effect predictions. A) SV eQTL classification performance for random forest classifiers trained on Borzoi outputs for 9,832 GTEx eQTL SVs using the default left-matched versus stitching strategies. Each point in the scatterplot represents a GTEx tissue. B) Predicted blood RNA-seq coverage for the reference sequence and alternative alleles for chr6:41,880,061 distal SNP (top) and chr6:41,897,089-41,897,626 (bottom) distal deletion. Both variants are associated with mean corpuscular hemoglobin and mean corpuscular volume, but they are in perfect LD (*r*^2^ = 1). C) ISM of the reference (top) and alternative deletion (bottom) alleles for the chr6:41,897,089-41,897,626 deletion variant.

### 2.6 Short tandem repeat variation effect prediction

Short tandem repeats (STRs) are prevalent in the human genome and often implicated in gene expression changes and disease [24]. However, tandem repeat variation is often excluded from genetic analyses due to bioinformatic challenges of genotyping such variants. As sequencing technologies and bioinformatic algorithms improve, it will be easier to estimate the contribution of tandem repeat variation to heritability of various traits. Meanwhile, it is useful to have a method to predict the effect of tandem repeat variation on expression. We assessed the ability of Borzoi with shift augmentation stitching to predict the effect of STRs with variable length on nearby gene expression using a set of fine-mapped STR eQTLs called from GTEx data [24]. Fig. 4A shows an example of predicted *NOP56* gene expression response to STR variation. The true effect size *β* in GTEx muscle has the same direction as the predicted slope *β*_*pred*_ from the ordinary least squares (OLS) regression fit, where we assumed linear dependency between allele number and *log*_2_ fold change (FC) of the allele expression relative to repeat number. We utilized the fine-mapped eQTL STRs from [24] with different PIP thresholds to ask whether variants with higher PIP, which are more likely to be causal, have more concordant prediction scores. For each likely causal STR eQTL, we fit an OLS on the dependency of the *log*_2_*FC* of a given gene expression versus variable STR count, filtered for regressions with a coefficient t-test *p <* 0.05, and compared the resulting slope *β*_*pred*_ with *β*_*true*_ (Methods). 36.1% (tissue-specific) and 36.6% (tissue-average) of the variants passed the significance test of the regression fit. Both the accuracy of predicted *β* direction and Spearman correlation of *β*_*pred*_ with *β*_*true*_ improve with higher PIP cutoff values (Fig. 4B,C). Taking the average of all GTEx RNA-seq tracks produced more concordant accuracies than matching tissue-specific tracks, perhaps because it helps denoise the predicted effect.

**Figure 4:**
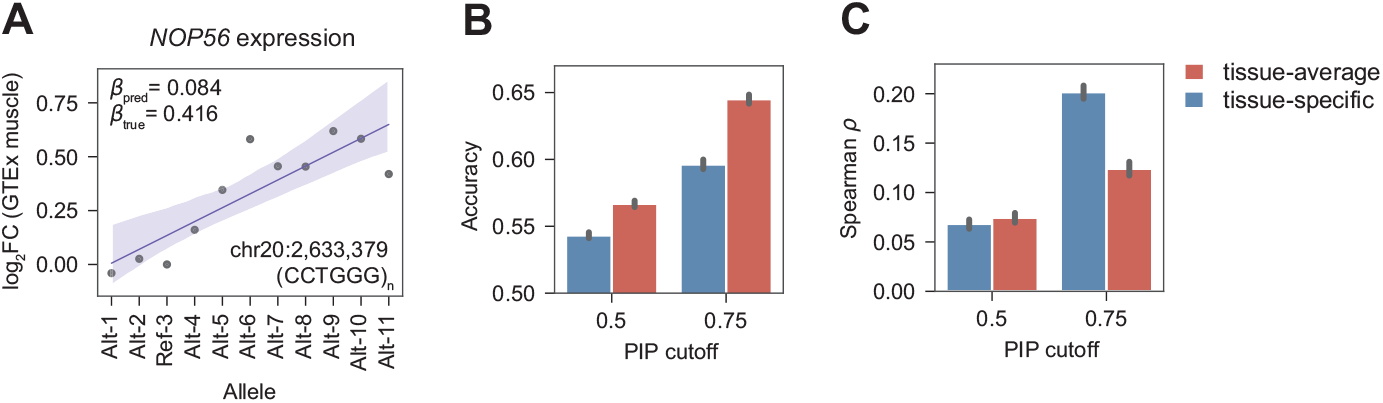
Shift augmentation stitching enables short tandem repeat variation effect prediction. A) Borzoi predicts *NOP56* expression is modulated by the expansion of short tandem motif (CCTGGG)_*n*_. *log*_2_*FC* of gene expression predicted in GTEx muscle tissue is shown on the yaxis with respect to the reference allele gene expression. The GRCh37/hg19 reference has 3 full repeats. B) Accuracy of sign prediction of the variable tandem repeat effect on gene expression *β* as a function of fine-mapped PIP threshold. Accuracy was computed using 1,000 bootstrap resampling experiments, with the standard deviation shown in the plot. For each PIP cutoff, we computed the accuracy using either predictions derived from averaging across all GTEx RNA-seq tracks or matching tissue-specific tracks only. C) Spearman correlation of predicted variable STR effect on gene expression *β* as a function of fine-mapped PIP threshold. Spearman *ρ* was computed using 1,000 bootstrap resampling experiments, with the standard deviation shown in the plot. For the three PIP thresholds, the numbers of variants with statistically significant (*p <* 0.05) regression fits are as follows: PIP=0.5: N=153 (tissue-average), N=179 (tissue-specific); PIP=0.75: N=69 (tissue-average), N=79 (tissue-specific).

### 2.7 *In silico* predictions of deletion effects are concordant with MPRA measurements

*In silico* mutagenesis (ISM) has emerged as a keystone regulatory sequence interpretation technique to identify the influential TF motifs and other sequence factors driving model predictions. Typical ISM mutates every reference nucleotide to its three alternatives, computing a prediction for each, and scoring the reference nucleotide based on the reference prediction relative to the average alternative. *In silico* deletion (ISD) of reference nucleotides could be an alternative to the typical ISM technique. We aimed to evaluate how well ISD works in comparison with ISM. Kircher et al. performed saturation mutagenesis of a set of disease-associated gene regulatory elements using massively parallel reporter assays (MPRA) [25]. Although they intended and focused on substitution mutations, error-prone PCR in the experimental protocol occasionally introduced deletions instead, which the authors measured and recorded. We sought to evaluate Borzoi ISD using a SUM score with shift augmentation stitching against these deletion measurements, as well as the substitution measurements and Borzoi ISM. We performed stitching of the reference sequence, as described above (visualization in Supplem. Fig. S1), requiring three forward passes through the model for this proof of concept; different designs could further reduce the computational expense if large-scale analysis were desired.

We selected two promoters with the highest correlation between the aggregated experimental substitution and deletion measurements for the same nucleotide positions: *HBG1* and *LDLR* (Pearson *r* = 0.502, 0.458 respectively). For substitutions, Borzoi ISM scores are concordant with the experimental substitution measurements (Fig. 5A,C,G,I) with Pearson *r* = 0.731 and 0.569 for *HBG1* and *LDLR*. Borzoi deletion scores are also concordant, albeit at lower levels, with both the experimental deletion (Pearson *r* = 0.219, 0.394; Fig. 5B,D,H,J) and substitution measurements (Pearson *r* = 0.284, 0.376; Fig. 5E,K). Borzoi substitution and deletion scores are correlated (Pearson *r* = 0.298, 0.609, Fig. 5F,L), with several interesting differences. For *HBG1*, the primary TF motifs emerge from both methods and the lower correlation is driven more by the flanking nucleotides. On the right, the T in the center of the G-rich motif is not sensitive to substitution, but Borzoi predicts its very sensitive to deletion, indicating that spacing may be critical. For LDLR, ISD misses an ATGACGT motif on the right that substitution predictions highlight (via Borzoi here and Enformer, too [1]), but the experimental data has no support for this motif, and it does not match anything in common TF databases well. For the majority of other tested promoters, we observe similar trends (Supplem. Fig. S2). This suggests that *in silico* deletion mutagenesis is a valid approach to probe regulatory sequence function.

**Figure 5:**
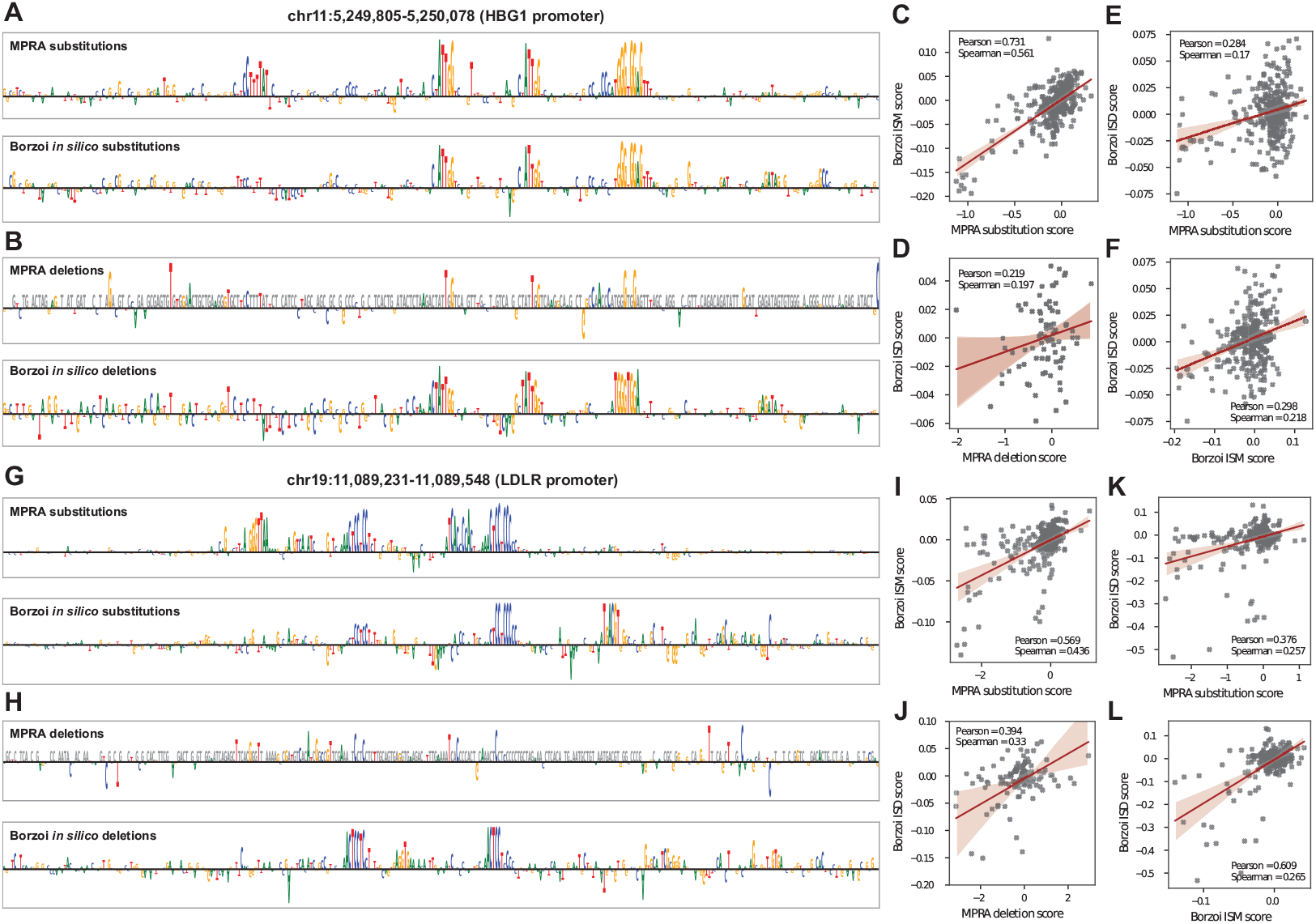
*In silico* nucleotide deletions are concordant with MPRA measurements. A), G) Comparison of MPRA saturation mutagenesis scores with Borzoi-derived ISM scores for *HBG1* and *LDLR* gene promoters. The nucleotide height quantifies the aggregate score for the reference nucleotide relative to the three alternative substitutions. B), H) Comparison of MPRA deletion scores with Borzoi-derived deletion scores for *HBG1* and *LDLR* gene promoters. The many missing experimental nucleotide deletions are shown in gray and excluded from the scatterplots. C), I) Scatterplots and correlations of Borzoi ISM scores with MPRA substitution measurements. D), J) Scatterplots and correlations of Borzoi deletion scores with MPRA deletion measurements. E), K) Scatterplots and correlations of Borzoi deletion scores with MPRA substitution measurements. F), L) Scatterplots and correlations of Borzoi deletion scores with Borzoi ISM scores.

## 3 Conclusions and future work

Dissecting the influence of genetic mutations on genes and their expression is a key task in genomics to harness and exploit the abundant genetic associations with traits. DNA sequence models like Enformer and Borzoi can predict genetic variant effects across assays and cell types to prioritize variants that affect transcript structure and expression level. However, indels are often omitted from variant prioritization analyses because of technical challenges. In this work, we demonstrated the presence of technical variance introduced by sequence shifts in sequence convolutional neural networks, exemplified by the Borzoi model. Even minor sequence shifts lead to misalignment of pooling block and output bin windows, resulting in prediction differences. In this work, we proposed and evaluated two shift mitigation strategies to reduce the influence of sequence shifts on indel effect prediction scores.

For models like Borzoi with sequential predictions in bins across the length of the sequence, the variance introduced by indels is best alleviated with a stitching strategy in which predictions from only the aligned regions of the sequence are extracted and concatenated to reconstruct the full prediction. To evaluate all strategies, we studied fine-mapped indel eQTLs from the GTEx project. We found that averaging the predictions from the left- and right-matched alternative allele sequences improves eQTL classification accuracy for both Borzoi and a single value prediction model, Basset. Stitching further improves accuracy for Borzoi, and represents our recommended strategy for future genetic analysis using this model. We further demonstrated the stitching strategy on larger structural variant and short tandem repeat eQTLs from GTEx, observing useful predictions on these more complex variants.

Shift augmentation stitching comes at the expense of excluding predictions in bins that overlap the indel sequence itself, which may not be appropriate in some cases. For example, as the indel size grows to be very large, predictions overlapping entire regulatory elements and/or exons may be disregarded. In this scenario, the included output bins will very likely also reflect substantial differences between the alternative and reference predictions, generating a large variant score anyway. Further, we expect that researchers will lean more heavily on prediction visualization strategies, rather than generic scores, as the indel size considered grows very large and the likelihood of molecular consequences saturates to certainty.

Despite a substantial reduction in the prediction variance introduced by indels with the stitching strategy, the distribution of predicted indel scores still diverges from the SNP score distribution. For applications that require fully matched distributions, additional techniques like quantile normalization, could be used. However, this may be counterproductive for larger indels, which likely have effect distributions reflecting larger influence. Fully alleviating the technical variance introduced by indels may require redesigning convolutional network architectures to remove all boundaries, e.g. by avoiding pooling and striding blocks and predicting at nucleotide resolution. The evaluations described here may be used to aid this architecture design evolution.

We additionally studied model sequence interpretation via nucleotide attribution scores derived from single nucleotide deletions, instead of the more common nucleotide substitutions of *in silico* mutagenesis (ISM) analysis. Regulatory sequence saturation deletion scores were concordant with experimental MPRA measurements for several human promoters with high quality data, and offered an intriguing alternative perspective for these regions. Thus, *in silico* deletion (ISD) analysis represents another option for ISM in cases, where deletions may represent the mutational process with higher fidelity.

In conclusion, our proposed strategies expand the applicability of regulatory sequence deep learning models to indel variants from small to large, including tandem repeats, enabling the comprehensive analysis of a wider range of genetic variation.

## 4 Acknowledgments

This work was funded by Calico Life Sciences LLC. The funder had no role in study design, data collection or analysis. Publication of the manuscript was approved after an internal scientific review process. We thank Johannes Linder, Han Yuan, Fanny Huang, and Madeleine Cule for helpful discussions and valuable feedback. We thank Luong Ruiz for help with code repositories and cloud computing infrastructure. The GTEx Project was supported by the Common Fund of the Office of the Director of the National Institutes of Health, with additional funds provided by the NCI, NHGRI, NHLBI, NIDA, NIMH and NINDS. The datasets used for the analyses described in this manuscript were obtained from dbGaP at http://www.ncbi.nlm.nih.gov/gap through dbGaP accession number phs000424.v9.p2.

## 5 Methods

### 5.1 DNA sequence deep learning architectures

Regulatory sequence models, such as Borzoi and Enformer, predict continuous coverage tracks from input DNA sequence of length *L*_*in*_. These architectures can be generalised as *f* (*X*) = *Y*, where 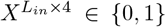 represents one-hot coded DNA sequence, and 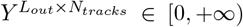,where *L*_*in*_ = 524, 288; *L*_*out*_ = 16, 384; *N*_*tracks*_ = 7, 611 for Borzoi. Alternatively, some architectures, such as Basset, predict a single value instead of a sequence, i.e. *L*_*out*_ = 1. The core elements of the architecture are convolutional blocks, interleaved with pooling layers that downsample incoming representations by computing a channel-wise summary statistic (such as maximum or mean) of a specified window, which will have width two throughout this analysis. When the input sequence shifts, pooling blocks will lead to differences in the extracted features, and the output bin boundaries will change.

### 5.2 Borzoi variant scores

We primarily study the recently published Borzoi model [2], which predicts 7,611 human ChIP, ATAC/DNase, CAGE, and RNA tracks via length 16,384 arrays where each position represents a 32 nt window. Briefly, the model consists of convolutional blocks, as describe above, to embed local sequence information, followed by transformer blocks that attend to longer sequence context and model distal interactions. The model has both human and mouse heads, but we only considered human tracks in this study.

To predict the effect of a variant on transcription and chromatin state, we center the input sequence at the variant and compute a forward pass through the model for the reference and alternative allele sequences. These forward passes include computing predictions for the forward and reverse complement sequences and averaging their outputs (after inverting the reverse complement transform). Generally, we then compute the inverse of any transformations performed on the original training data and then compute *log*_2_(*y* + 1) on the bin values.

Finally, we seek to collapse the length axis in order to produce a single score for each output track. To consider the signed effect (e.g. up- or down-regulating gene expression), we compute the ‘SUM’ score by summing values across the length axes and taking the difference between alleles: 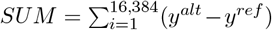.To consider a more sensitive, but unsigned effect, we compute the ‘L2’ score as an L2 norm of the difference between length-wise vectors: 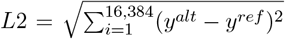. We compute these scores using four model replicates and take their average as an ensemble.

### 5.3 Indel shift augmentation strategies

While SNPs involve a simple substitution of one nucleotide for another, insertions and deletions alter the sequence more dramatically, from the perspective of a convolutional neural network. Using standard VCF coding, the most natural insertion would add the new nucleotides to the right of center and push several nucleotides (determined by the insertion length) off the right edge of the sequence (Fig. 2A, Supplem. Fig. S1). Because the left side of the input sequence remains unchanged relative to the reference sequence, we refer to this method as “left-matched”. However, a symmetric “right-matched” shift, in which the new nucleotides are added to the left of the center and several nucleotides are pushed off the left edge of the sequence, is equally valid. Analogously, a natural deletion would remove nucleotides to the right of center and introduce padded sequence at the right edge of the sequence (“left-matched”) or remove nucleotides to the left of center and introduce padded sequence at the left edge of the sequence (“right-matched”).

We propose computing both of these versions for each indel, since they will produce different but equally valid scores. Subsequently, they must be combined into a single vector of scores (one per track). In our simplest strategy, we treat these as an ensemble and take their average. When *L*_*out*_ *>* 1, an intriguing alternative emerges. Because the “left-matched” and “right-matched” versions each match the reference on one side of the variant, the two halves can be stitched together into a prediction that matches the reference across the full sequence. Specifically, we split the prediction pairs exactly at the center of the length axis and concatenate the left-matched predictions to the right-matched predictions to form a full prediction tensor. We disregard the right portion of the left-matched predictions and the left portion of the right-matched predictions. We then compute variant scores comparing the reference predictions to these stitched alternative predictions.

Upon close inspection, we realized that computing deletion predictions as described above does not exactly produce aligned predictions and double counts the nucleotides to the left of the deletion. The proper way to align the predictions is to perform the shifts to the sequence with the added nucleotides. That is, deletions should be treated as insertions to the reference, and the shift recipe described above should be applied to the reference, instead of the alternative sequence. See Supplementary Figure S1 for visualization representations of these scenarios.

In all analyses here, we center the variant, but some cases might prefer to relax this requirement. For example, large-scale SNP scoring can be accelerated by predicting the reference sequence once, followed by predicting alternative sequences for all variants that overlap the reference sequence. With large sequences containing many variants, the computation time will approach 2x faster by avoiding redundant reference sequence predictions. However, variants can occur at arbitrary locations in the sequence, which challenges the precision of the stitching strategy. Nevertheless, one can adopt the strategy despite the imprecision, and a minimum of one output bin will contain some degree of misalignment.

These indel scoring strategies and example visualization notebooks are implemented in the open source baskerville (https://github.com/calico/baskerville) and Borzoi (https://github.com/calico/borzoi) code repositories.

### 5.4 Basset variant scores

To explore shift strategies for models with *L*_*out*_ = 1, such as binary peak predictors, we chose the Basset model [21]. Basset predicts 164 DNase peak probabilities for a 600 nt input sequence, using convolution/pooling blocks that operate at window widths of 3, 4, and 4. As described above, one can apply left-matched or right-matched shifts to predict alternative indel alleles. We computed Basset predictions using the Kipoi model repository [26]. Input sequences for Basset were centered at the indel of interest, and we compared reference and alternative allele predictions using subtraction: *y*^*alt*^ − *y*^*ref*^ .

### 5.5 Fine-mapped eQTL data and discrimination task

We made use of GTEx v8 eQTLs fine-mapped by the SuSiE algorithm [27]. This set contains 1,168 unique indels up to size 8 from 49 tissues with a credible set posterior inclusion probability (PIP)*>*0.9. The first benchmark is to discriminate between the causal (positive) eQTLs and negative variants matched for indel size and PIP*<*0.01 but |Z-score| *>* 4 for the same gene. If unavailable for a given gene, we selected from the genome-wide set with PIP*<*0.01 and |Z-score| *>* 6. These variants, and thus our predictions, are mapped to the GRCh38/hg38 human genome assembly. For this analysis, we focused on L2 scores, which were shown to be more informative for eQTL classification [2]. For each tissue, we trained a random forest classifier using RandomForestClassifier from sklearn.ensemble with the following parameters: min samples leaf=1, max depth=64, n estimators=100, max features=“log2”. We performed 8 fold cross-validation and report the mean AUROC of 100 independent stochastic classifier training procedures. This evaluation considers the full richness of the prediction vector across Borzoi and Basset tracks to identify eQTL indels.

Second, we evaluated how well the indel effect size on target gene expression can be estimated with Borzoi predictions. For this analysis, we sliced the matching RNA-seq tracks for each GTEx tissue, computed the SUM metric, and averaged across the sliced tracks. We made use of the SuSiE *β* posterior statistic, and filtered out all eQTLs that are significant for multiple genes, but influence gene expression of the genes in opposing directions. For the eQTLs with consistent effect directions, we took the mean of the *β* posterior if this eQTL affected multiple genes as the true effect size. For tissues with *>* 35 consistent fine-mapped indel eQTLs (24 tissues), we computed the Spearman *ρ* correlation between SUM scores and eQTL coefficients.

### 5.6 Structural variant eQTL classification benchmark

GTEx structural variant (SV) eQTLs were independently retrieved from analysis performed by Kirsche et al. with the Jasmine and Iris tools [22]. We focused on lead SV eQTLs for each gene-tissue pair that passed the FDR threshold and limited SV size to 1/16th of the input sequence size (524288*/*16 = 32768 bp) to ensure sufficient sequence context around the SV. For variants that had marginally significant associations with multiple genes, we only considered one eGene with the lowest association p-value, producing a total set of 9,832 unique variants. To create a negative set for a classification task, we retrieved all variants from the cohort-level call set that did not have a marginal eQTL association. We filtered away variants with AF≤0.05. We matched each positive SV to a negative SV with the closest available size, for which at least one expressed gene exon fell within the input sequence. We made use of gene expression from the GTEx portal https://www.gtexportal.org/home/downloads/adult-gtex/bulk_tissue_expression: GTEx Analysis 2017-06-05 v8 RNASeQCv1.1.9 gene median tpm.gct. These variants, and thus our predictions, are mapped to the GRCh38/hg38 human genome assembly. Finally, we trained random forest classifiers on Borzoi L2 scores, using the stitch strategy, with the procedure described above.

### 5.7 Tandem repeat eQTL *β* prediction benchmark

GTEx short tandem repeat (STR) eQTLs were independently retrieved from the Supplementary Data of [24]. We only considered variant-gene pairs that had the same effect direction on gene expression in all significant tissues. We note that the number of repeats found by genotyping in the original study does not always coincide with the number of repeats counted from the reference genome assembly. For each variant, we found the reference repeat number by counting the number of full repeats in the reference genome (GRCh37/hg19, as in the original study), and iteratively removed up to 4 repeats (or the number of repeats corresponding to reference TR, if less than 4), and added up to 4 repeats, and predicted resulting gene expression using Borzoi with shift augmentation stitching. If the STR started with a partial motif, the repeats were removed or appended from the first found full repeat coordinate. We then fitted a linear regression model on gene expression *log*_2_*FC* with respect to the reference allele versus allele number (absolute number of repeats) using ordinary least squares from python statsmodels package. We used either the average of all available GTEx tissue RNA-seq tracks, or only tissue-matched GTEx RNA-seq tracks. Unlike most variant scoring, the regression statistics offer a built-in confidence measure, and we only included variants with significant regression coefficient p-value (t-test *p <* 0.05) in the analysis. We then used different PIP thresholds to compute the accuracy of direction of change with STR expansion, and Spearman correlation of the fitted *β*_*pred*_ with fine-mapped *β*_*true*_ reported in the original study.

### 5.8 *in silico* versus MPRA substitution and deletion mutagenesis

Borzoi-generated *in silico* substitution mutagenesis scores (ISM) for the MPRA saturation mutagenesis dataset [25] were computed as follows. For each position, we computed predictions for the reference sequence and the three substitution variants, with the input sequence centered on each variant. We cropped the output window to the center 4kb to capture local changes for promoters. Fold change scores were then computed gene-agnostically for DNase and H3K4me3 tracks, while RNA tracks were computed gene-specifically as: 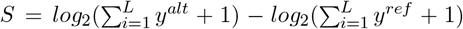, where *L* = 4*kb* or *L* = *bins* ∩ *exons*, respectively. The total score for a variant is the average of all cell type-matched DNase, H3K4me3 and RNA track scores. To assign each position a single score, we computed the negative mean of the three mutation scores per position; i.e. if mutating away from a given nucleotide decreases the coverage of the tracks, it will have a positive ISM score.

For in silico deletion mutagenesis (ISD), we computed scores similarly, but using one-nucleotide deletions instead of substitutions. As described above for general deletion variants, we applied reference sequence stitching for ISD calculations. We made use of the same cell type to promoter matching as in [2]. The experimental MPRA score was derived by averaging the *log*_2_ variant expression effect values from all three alternative alleles [25]. For visualization, reference nucleotides are shown with the respective ISM or ISD RNA-only scores. Deletion scores missing in experimental data were omitted from the correlation analysis.

Although reference-stitching maintains precise alignment, it costs three forward passes per nucleotide scored. For large-scale ISD analysis, one could relax this constraint using the follow strategy that costs a single forward pass per nucleotide. Arrange the sequence so that the output bins of interest are safely to the right of the center, and compute the reference sequence prediction once.

For each nucleotide, introduce the deletion using a right-matched strategy so that the output bins of interest are unaffected and comparison is clean.

## A Supplementary figures

**Figure S1:**
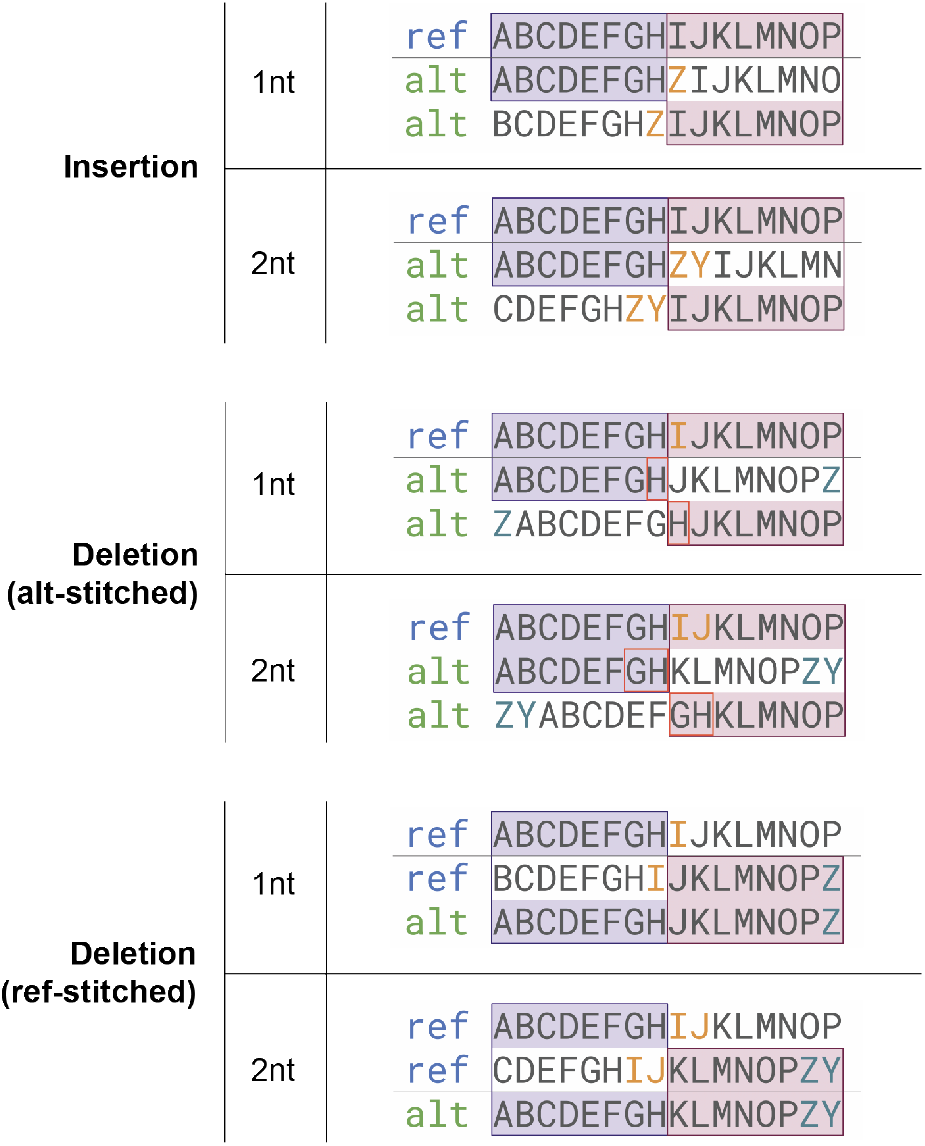
Shift augmentation stitching visualization for insertions and deletions. Inserted or deleted nucleotides are shown in orange, while nucleotides compensating for the deletion on the edge of the sequence are shown in teal. Purple and pink boxes show the left- and right-matched stitched parts of sequence, respectively. For insertions, the alternative alleles shifted to the left and to the right by the number of inserted nucleotides are stitched to avoid mismatched nucleotides, whereas for deletions the reference is shifted by *N* = *len*(*deletion*) to the left, and stitched before computing the score with the alternative. Originally, we applied alternative allele stitching for both insertion and deletion, but upon careful examination we noticed that with alt-stitched strategy, deletions will have repeated nucleotides at the stitch boundary (see middle diagram, repeated nucleotides are highlighted in red boxes). Performing the stitching strategy to the reference instead maintains precise alignments and avoids this double counting

**Figure S2:**
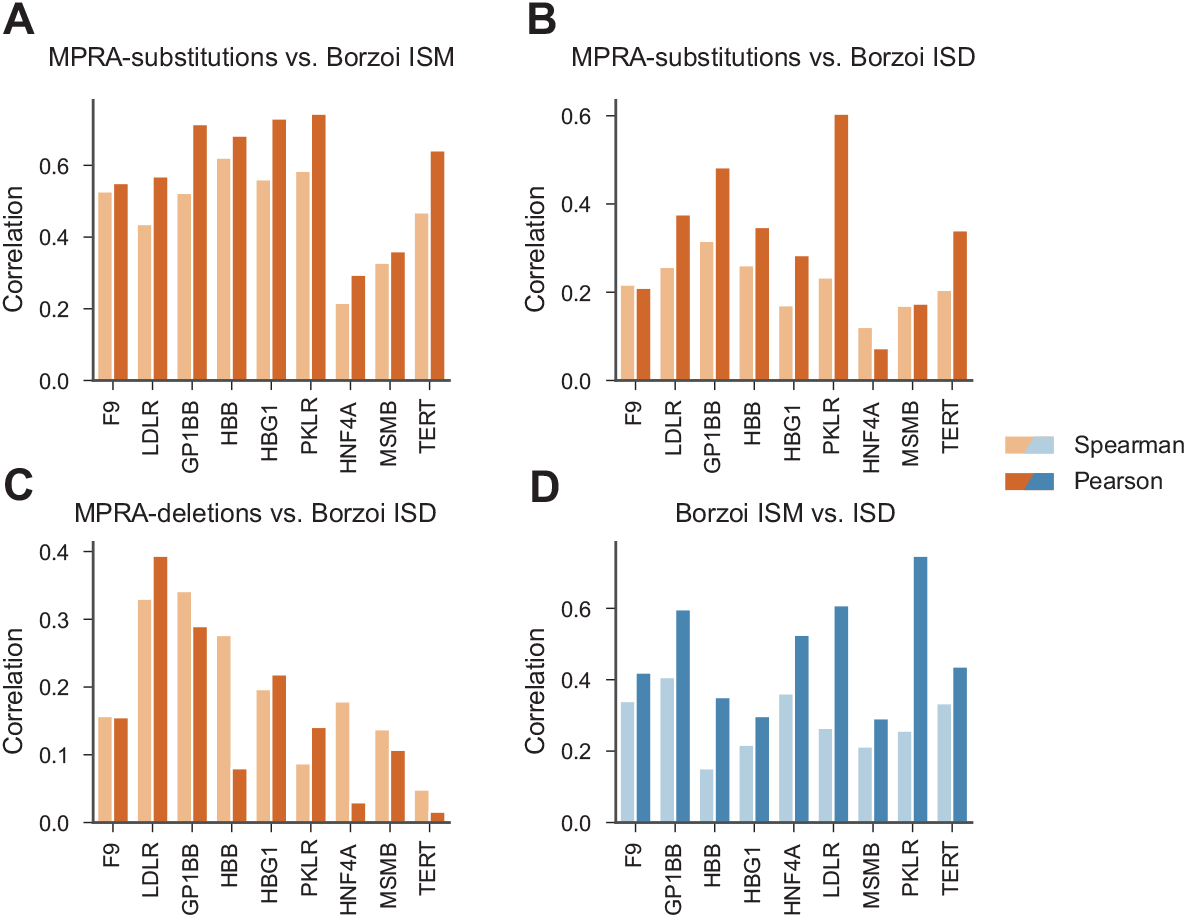
Associations between Borzoi-derived ISM and ISD scores versus MPRA saturation mutagenesis scores for gene promoters from [25]. A) Correlations between MPRA substitution scores and Borzoi ISM. B) Correlations between MPRA substitution scores and Borzoi ISD. C) Correlations between MPRA deletion scores and Borzoi ISD. D) Concordance between Borzoi ISM and ISD scores for the same nucleotide positions. Spearman and Pearson correlations are shown in each plot for all experimentally available nucleotide positions.

## References

[1] Žiga Avsec, Vikram Agarwal, Daniel Visentin, Joseph R Ledsam, Agnieszka Grabska-Barwinska, Kyle R Taylor, Yannis Assael, John Jumper, Pushmeet Kohli, and David R Kelley. Effective gene expression prediction from sequence by integrating long-range interactions. Nature methods, 18(10):1196–1203, 2021.

[2] Johannes Linder, Divyanshi Srivastava, Han Yuan, Vikram Agarwal, and David R. Kelley. Predicting RNA-seq coverage from DNA sequence as a unifying model of gene regulation. Nat. Genet., pages 1–13, January 2025.

[3] Kishore Jaganathan, Sofia Kyriazopoulou Panagiotopoulou, Jeremy F McRae, Siavash Fazel Darbandi, David Knowles, Yang I Li, Jack A Kosmicki, Juan Arbelaez, Wenwu Cui, Grace B Schwartz, et al. Predicting splicing from primary sequence with deep learning. Cell, 176(3):535– 548, 2019.

[4] Tony Zeng and Yang I Li. Predicting rna splicing from dna sequence using pangolin. Genome biology, 23(1):1–18, 2022.

[5] Johannes Linder, Samantha E Koplik, Anshul Kundaje, and Georg Seelig. Deciphering the impact of genetic variation on human polyadenylation using aparent2. Genome Biology, 23(1):1– 33, 2022.

[6] Adam Y. He and Charles G. Danko. Dissection of core promoter syntax through single nucleotide resolution modeling of transcription initiation. bioRxiv, page 2024.03.13.583868, April 2024.

[7] Kelly Cochran, Melody Yin, Anika Mantripragada, Jacob Schreiber, Georgi K. Marinov, and Anshul Kundaje. Dissecting the cis-regulatory syntax of transcription initiation with deep learning. bioRxiv, page 2024.05.28.596138, June 2024.

[8] Kseniia Dudnyk, Donghong Cai, Chenlai Shi, Jian Xu, and Jian Zhou. Sequence basis of transcription initiation in the human genome. Science, 384(6694), April 2024.

[9] Vikram Agarwal and David R Kelley. The genetic and biochemical determinants of mrna degradation rates in mammals. Genome Biology, 23(1):245, 2022.

[10] Kathleen M. Chen, Aaron K. Wong, Olga G. Troyanskaya, and Jian Zhou. A sequence-based global map of regulatory activity for deciphering human genetics. Nat. Genet., 54:940–949, July 2022.

[11] Žiga Avsec, Melanie Weilert, Avanti Shrikumar, Sabrina Krueger, Amr Alexandari, Khyati Dalal, Robin Fropf, Charles McAnany, Julien Gagneur, Anshul Kundaje, and Julia Zeitlinger. Base-resolution models of transcription-factor binding reveal soft motif syntax. Nat. Genet., 53:354–366, March 2021.

[12] Geoff Fudenberg, David R. Kelley, and Katherine S. Pollard. Predicting 3D genome folding from DNA sequence with Akita. Nat. Methods, 17:1111–1117, November 2020.

[13] Jian Zhou. Sequence-based modeling of three-dimensional genome architecture from kilobase to chromosome scale. Nat. Genet., 54:725–734, May 2022.

[14] Jian Zhou, Christopher Y Park, Chandra L Theesfeld, Aaron K Wong, Yuan Yuan, Claudia Scheckel, John J Fak, Julien Funk, Kevin Yao, Yoko Tajima, et al. Whole-genome deep-learning analysis identifies contribution of noncoding mutations to autism risk. Nature genetics, 51(6):973–980, 2019.

[15] Tabassum Fabiha, Ivy Evergreen, Soumya Kundu, Anusri Pampari, Sergey Abramov, Alexandr Boytsov, Kari Strouse, Katherine Dura, Weixiang Fang, Gaspard Kerner, et al. A consensus variant-to-function score to functionally prioritize variants for disease. bioRxiv, 2024.

[16] Siwei Chen, Laurent C. Francioli, Julia K. Goodrich, Ryan L. Collins, Masahiro Kanai, Qingbo Wang, Jessica Alföldi, Nicholas A. Watts, Christopher Vittal, Laura D. Gauthier, Timothy Poterba, Michael W. Wilson, Yekaterina Tarasova, William Phu, Riley Grant, Mary T. Yohannes, Zan Koenig, Yossi Farjoun, Eric Banks, Stacey Donnelly, Stacey Gabriel, Namrata Gupta, Steven Ferriera, Charlotte Tolonen, Sam Novod, Louis Bergelson, David Roazen, Valentin Ruano-Rubio, Miguel Covarrubias, Christopher Llanwarne, Nikelle Petrillo, Gordon Wade, Thibault Jeandet, Ruchi Munshi, Kathleen Tibbetts, Anne O’Donnell-Luria, Matthew Solomonson, Cotton Seed, Alicia R. Martin, Michael E. Talkowski, Heidi L. Rehm, Mark J. Daly, Grace Tiao, Benjamin M. Neale, Daniel G. MacArthur, and Konrad J. Karczewski. A genomic mutational constraint map using variation in 76,156 human genomes. Nature, 625:92– 100, January 2024.

[17] Sarah A. Gagliano, Sebanti Sengupta, Carlo Sidore, Andrea Maschio, Francesco Cucca, David Schlessinger, and Gonçalo R. Abecasis. Relative impact of indels versus SNPs on complex disease. Genet. Epidemiol., 43(1):112, November 2018.

[18] Matthew T. Weirauch, Ally Yang, Mihai Albu, Atina G. Cote, Alejandro Montenegro-Montero, Philipp Drewe, Hamed S. Najafabadi, Samuel A. Lambert, Ishminder Mann, Kate Cook, Hong Zheng, Alejandra Goity, Harm van Bakel, Jean-Claude Lozano, Mary Galli, Mathew G. Lewsey, Eryong Huang, Tuhin Mukherjee, Xiaoting Chen, John S. Reece-Hoyes, Sridhar Govindarajan, Gad Shaulsky, Albertha J. M. Walhout, François-Yves Bouget, Gunnar Ratsch, Luis F. Larrondo, Joseph R. Ecker, and Timothy R. Hughes. Determination and inference of eukaryotic transcription factor sequence specificity. Cell, 158(6):1431–1443, September 2014.

[19] G. Mavrothalassitis and J. Ghysdael. Proteins of the ETS family with transcriptional repressor activity. Oncogene, 19(55):6524–6532, December 2000.

[20] Michel Fausther, Elise G. Lavoie, Jessica R. Goree, and Jonathan A. Dranoff. An Elf2-like transcription factor acts as repressor of the mouse ecto-5′-nucleotidase gene expression in hepatic myofibroblasts. Purinergic Signalling, 13(4):417, December 2017.

[21] David R Kelley, Jasper Snoek, and John L Rinn. Basset: learning the regulatory code of the accessible genome with deep convolutional neural networks. Genome research, 26(7):990–999, 2016.

[22] Melanie Kirsche, Gautam Prabhu, Rachel Sherman, Bohan Ni, Alexis Battle, Sergey Aganezov, and Michael C. Schatz. Jasmine and Iris: population-scale structural variant comparison and analysis. Nat. Methods, 20:408–417, March 2023.

[23] Marsha M. Wheeler, Adrienne M. Stilp, Shuquan Rao, Bjarni V. Halldórsson, Doruk Beyter, Jia Wen, Anna V. Mihkaylova, Caitlin P. McHugh, John Lane, Min-Zhi Jiang, Laura M. Raffield, Goo Jun, Fritz J. Sedlazeck, Ginger Metcalf, Yao Yao, Joshua B. Bis, Nathalie Chami, Paul S. de Vries, Pinkal Desai, James S. Floyd, Yan Gao, Kai Kammers, Wonji Kim, Jee-Young Moon, Aakrosh Ratan, Lisa R. Yanek, Laura Almasy, Lewis C. Becker, John Blangero, Michael H. Cho, Joanne E. Curran, Myriam Fornage, Robert C. Kaplan, Joshua P. Lewis, Ruth J. F. Loos, Braxton D. Mitchell, Alanna C. Morrison, Michael Preuss, Bruce M. Psaty, Stephen S. Rich, Jerome I. Rotter, Hua Tang, Russell P. Tracy, Eric Boerwinkle, Goncalo R. Abecasis, Thomas W. Blackwell, Albert V. Smith, Andrew D. Johnson, Rasika A. Mathias, Deborah A. Nickerson, Matthew P. Conomos, Yun Li, Unnur orsteinsdóttir, Magnuś K. Magnuśson, Kari Stefansson, Nathan D. Pankratz, Daniel E. Bauer, Paul L. Auer, and Alex P. Reiner. Whole genome sequencing identifies structural variants contributing to hematologic traits in the NHLBI TOPMed program. Nat. Commun., 13(7592):1–18, December 2022.

[24] Stephanie Feupe Fotsing, Jonathan Margoliash, Catherine Wang, Shubham Saini, Richard Yanicky, Sharona Shleizer-Burko, Alon Goren, and Melissa Gymrek. The impact of short tandem repeat variation on gene expression. Nat. Genet., 51:1652–1659, November 2019.

[25] Martin Kircher, Chenling Xiong, Beth Martin, Max Schubach, Fumitaka Inoue, Robert J. A. Bell, Joseph F. Costello, Jay Shendure, and Nadav Ahituv. Saturation mutagenesis of twenty disease-associated regulatory elements at single base-pair resolution. Nat. Commun., 10(3583):1–15, August 2019.

[26] Žiga Avsec, Roman Kreuzhuber, Johnny Israeli, Nancy Xu, Jun Cheng, Avanti Shrikumar, Abhimanyu Banerjee, Daniel S. Kim, Thorsten Beier, Lara Urban, Anshul Kundaje, Oliver Stegle, and Julien Gagneur. The Kipoi repository accelerates community exchange and reuse of predictive models for genomics. Nat. Biotechnol., 37:592–600, June 2019.

[27] Qingbo S Wang, David R Kelley, Jacob Ulirsch, Masahiro Kanai, Shuvom Sadhuka, Ran Cui, Carlos Albors, Nathan Cheng, Yukinori Okada, et al. Leveraging supervised learning for functionally informed fine-mapping of cis-eqtls identifies an additional 20,913 putative causal eqtls. Nature Communications, 12(1):3394, 2021.

